# Simultaneous development and periodic clustering of simple and complex cells in visual cortex

**DOI:** 10.1101/782151

**Authors:** Gwangsu Kim, Jaeson Jang, Se-Bum Paik

## Abstract

Neurons in the primary visual cortex (V1) are often classified as simple or complex cells, but it is debated whether they are discrete hierarchical classes of neurons developing sequentially, or if they represent a continuum of variation within a single class of cells developing simultaneously. Herein, we show that simple and complex cells may arise simultaneously from the universal process of retinal development. From analysis of the cortical receptive fields in cats, we show evidence that simple and complex cells originate from the periodic variation of ON-OFF segregation in the feedforward projection of retinal mosaics, by which they organize into periodic clusters in V1. Our key prediction that clusters of simple and complex cells correlate topographically with orientation maps was confirmed by data in cats. Our results suggest that simple and complex cells are not two distinct neural populations but arise from common retinal afferents, simultaneous with orientation tuning.

**Highlights:**

- Simple and complex cells arise simultaneously from retinal afferents.
- Simple/complex cells are organized into periodic clusters across visual cortex.
- Simple/complex clusters are topographically correlated with orientation maps.
- Development of clustered cells in V1 is explained by the Paik-Ringach model.

## Introduction

Neurons in the primary visual cortex (V1) are often classified as simple or complex cells^1^ by their characteristic organization of spatial receptive fields and the temporal dynamics of their response to stimuli. In traditional classifications, simple cells have segregated ON/OFF sub-regions of receptive fields and generate highly modulated sinusoidal response (*F*_*1*_*/F*_*0*_ > 1) to drifting gratings stimuli, while complex cells have largely overlapping ON/OFF sub-regions and generate weak modulation of response (*F*_*1*_*/F*_*0*_ < 1, Fig. 1a)^1–7^. As suggested in the pioneering study of Hubel and Wiesel, simple and complex cells have often been considered to imply a hierarchically distinct functional architecture for visual processing^1,8–11^, so that simple cells pool thalamic inputs^12,13^, while complex cells then pool inputs from the simple cells (Fig. 1b)^14,15^.

**Figure 1.**
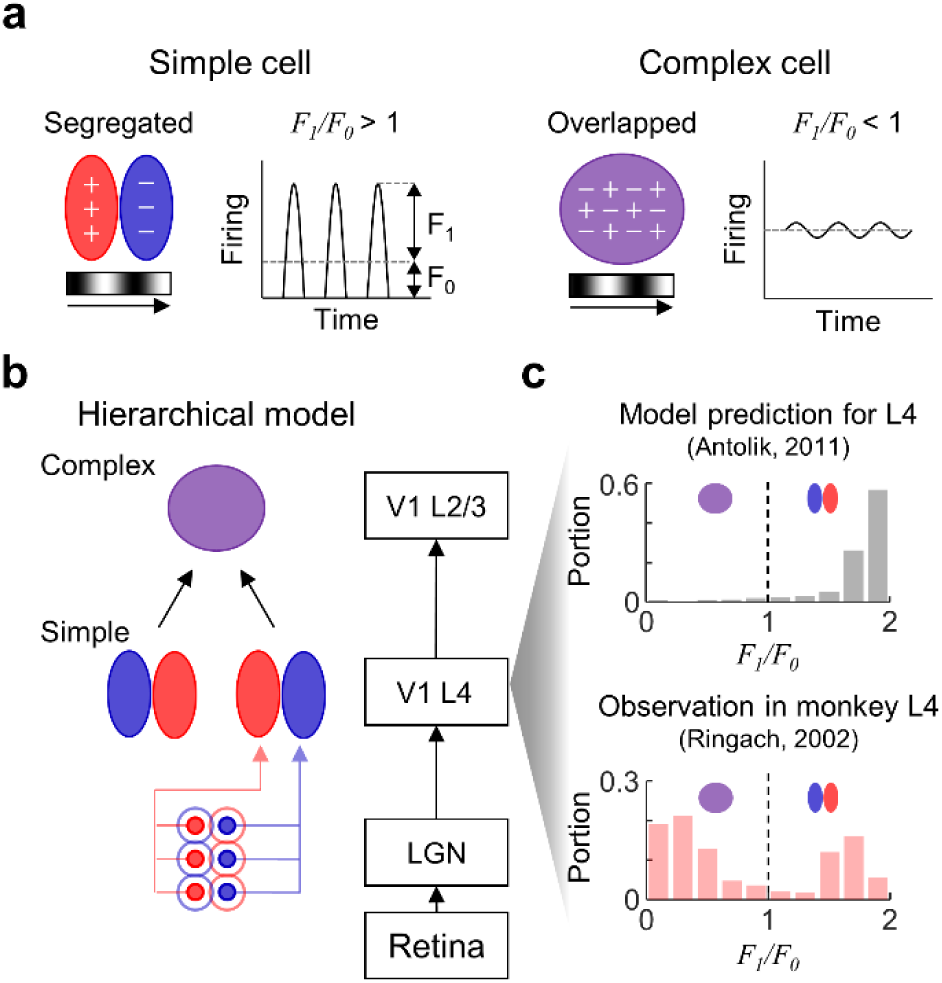
Parallel development of simple and complex cells in the primary visual cortex. (a) Illustration of simple (left) and complex (right) cells. Receptive fields and response profiles to a drifting sinusoidal grating stimulus are described. Red area (+): ON subregion. Blue area (−): OFF subregion. Purple area (+&−): both ON and OFF subregion. *F*_*1*_*/F*_*0*_: ratio of 1^st^ harmonic amplitude to mean elevation of firing rate. (b) Schematic of the classical hierarchical model. (c) Bimodal histograms of *F*_*1*_*/F*_*0*_ in Layer 4 of adult monkeys (red bar, adapted from reference^16^) and prediction of hierarchical model (gray bar, adapted from reference^10^).

Although the conventional hierarchical model predicts that neurons in the early layer are mostly simple cells^1,10^ (Fig. 1c, top), it was observed that complex cells coexist with simple cells in layer 4 of monkey V1^16^, the earliest cortical stage that receives direct feedforward inputs from the thalamus (Fig. 1c, bottom; see Supplementary Fig. 2 for cats^9^ and tree shrews^17^), implying that simple and complex cells may arise simultaneously from a common origin. It was also reported that simple- and complex-like receptive fields can arise together in the primary auditory cortex, when retinal afferents are rewired to give inputs to the auditory thalamus^18^. This result suggests that feedforward afferents can induce both simple and complex cells from common retinal afferents.

Furthermore, subsequent studies have raised the possibility that simple and complex neurons are not clearly distinct populations but might be variations within a continuous spectrum^19–21^. Experimental evidence supporting this notion has been reported—conventional criteria for distinguishing simple and complex cells are susceptible to stimulus modulation^22–24^ and nonlinearity of the spike threshold might be a prime determinant for simple and complex classes^25^. Thus, these results raise questions on the origin of simple and complex cells: Might simple and complex cells arise from non-distinctive neural circuits? If so, then what possible mechanism is there for the development of such a functional variation?

Recent studies on the retinal origin of cortical tunings provide clues regarding the wiring of simple and complex tunings in the early layer of visual cortex^26–28^. A number of studies have reported evidence to strengthen the role of ON/OFF retinal afferents in developing diverse functional tuning of neurons in V1. For example, the orientation preference of a cortical column can be predicted by the local arrangement of ON/OFF afferents^29,30^ and other functional tunings such as direction selectivity^31^, ON/OFF polarity, and ocular dominance^32^ are observed to develop from the integration of thalamic inputs. Considering that local thalamic receptive fields preserve those of retinal ganglion cells (RGC), all these results support the notion that the spatial distribution of ON/OFF receptive fields in retinal mosaics determine the formation of orientation tuning and their topographic organization^26,27,33–35^.

Here, we propose that the simple and complex tuning of V1 neurons arises from the periodic variation of a common retinal mosaics structure, topographically correlated with the orientation tuning of underlying neurons. From the analysis of data in cats^32^, we show evidence that neuronal variation from simple to complex cells can be predicted from the segregation between local ON and OFF feedforward afferents. Importantly, systematic formation of distinct clusters of simple and of complex cells was observed across V1, the spatial period of which was matched to that of underlying orientation maps. We also show that the Paik-Ringach model^26,27^ provides a plausible developmental mechanism for the observed results, implying that simple/complex tuning and orientation selectivity may have a common origin. Our further prediction that pinwheels on the orientation map and clusters of simple/complex tuning are topographically correlated, was validated by the analysis of cat data.

Overall, our findings suggest that simple and complex cells in V1 develop simultaneously from structured inputs from the retina, which enables a parallel architecture of the simple and complex tuning in V1 that is tightly correlated with the topography of other functional maps.

## Results

### Simple and complex cells from the spatial arrangement of ON/OFF retinal afferents

Based on the theory that functional tuning in the visual cortex originates from the afferent of ON and OFF RGC mosaics^26,27,33–35^, we hypothesized that both simple and complex cells in V1 are initially seeded by the local projection of feedforward afferents, and that the variation of cell types in development is dependent on the spatial distribution of ON and OFF receptive fields imprinted in RGC mosaics (Fig. 2a). We introduced our model idea by investigating the profile of retinal mosaics data of ON-center and OFF-center receptive fields (RFs, Fig. 2b)^36^. As previously reported^36,37^, the nearest neighbor distance between different types of RF centers (*d*_*ON-OFF*_) appeared smaller than that between the same type (average of *d*_*ON-ON*_ = 116 μm and *d*_*OFF-OFF*_ = 106 μm), thus the nearest neighbor of an ON cell appears to be an OFF cell, and vice versa. The profile of this ON-OFF distance (*d*_*ON-OFF*_) measured from RGC mosaics data^36^ showed a wide variation, well fitted to a Gaussian distribution (mean = 56.4 μm, standard deviation = 14.3 μm, *R*^*2*^ = 0.91) (Fig. 2b, bottom histogram).

**Figure 2.**
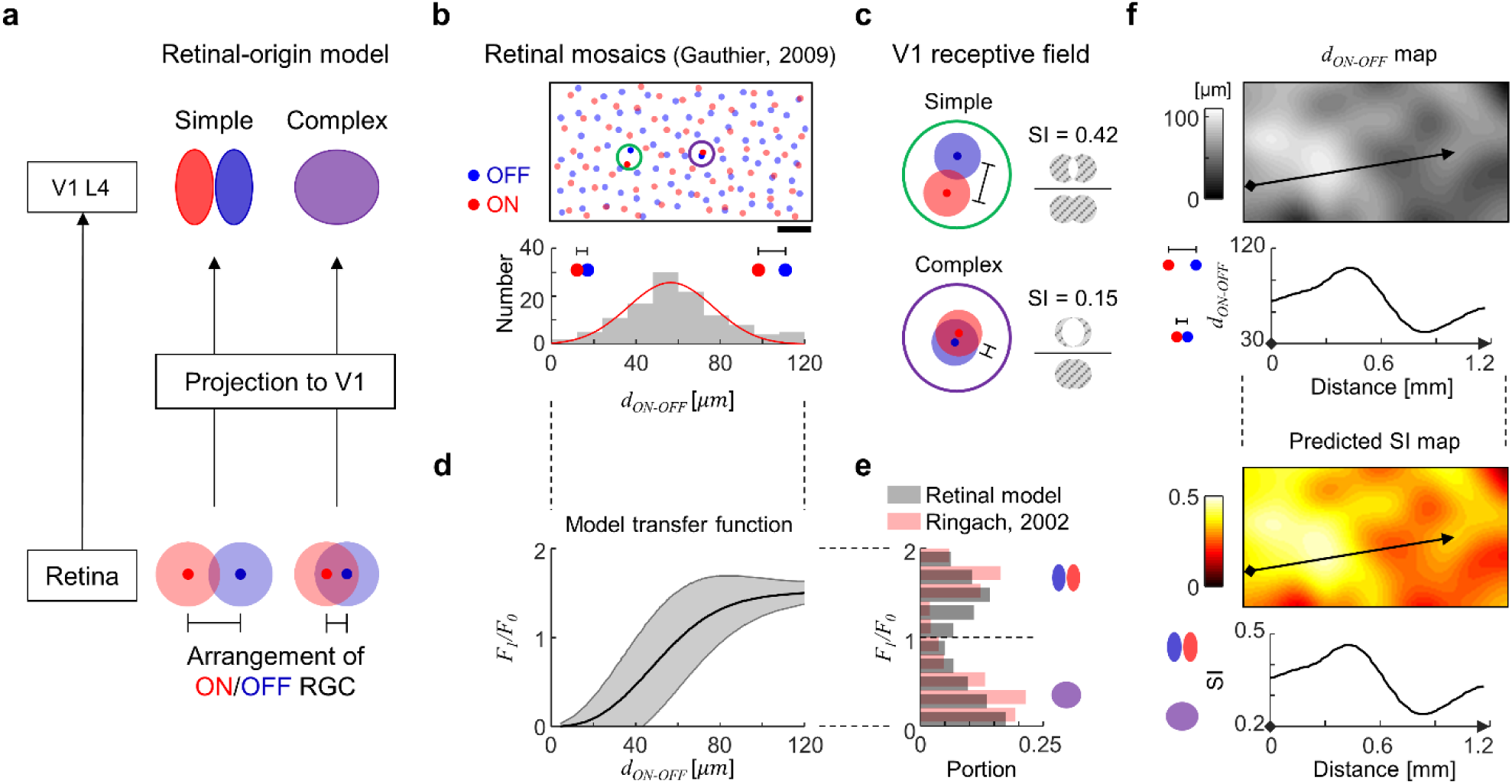
Retinal-origin model of simple and complex cells. (a) Schematic of retinal-origin model. ON- and OFF-center RGC receptive fields are represented as red and blue circles (dots represent the center of mass). (b) ON and OFF-center RGC receptive field mosaic data from monkeys^36^. Scale bar: 0.2 mm. ON-OFF dipoles (N = 116) were defined as a line from each OFF cell to the nearest ON cell, and *d*_*ON-OFF*_ denotes the size of the dipole. Bottom histogram represents the distribution of *d*_*ON-OFF*_, fitted to a Gaussian (Red curve). (c) Example receptive fields of local ON and OFF RGC afferents (from green and purple circles in b). Simpleness index (SI) measures the spatial segregation between ON and OFF receptive field sub-regions. (d) A nonlinear transformation between *F*_*1*_*/F*_*0*_ and ON-OFF distance obtained by an analytic model (see Supplementary Information). Shaded area represents standard deviation. (e) Resulting bimodal distribution of *F*_*1*_*/F*_*0*_ predicted by the model (gray, Hartigan’s dip test, *p* < 10^−5^) is compared with the histogram of *F*_*1*_*/F*_*0*_ in Figure 1(b) (red). (f) Spatial map and example 1-D profile (along black arrows) of *d*_*ON-OFF*_ obtained from mosaic data in (a) (Top) and those of SI (Bottom).

Our main hypothesis is that this spatial organization of ON and OFF RGC can constrain the tuning of the connected V1 neurons as either simple or complex cells, via statistical wiring from the retina to V1^26,27,33–35^. When the distance between ON and OFF RGC is large (Fig. 2c, green circle, *d*_*ON-OFF*_ = 87 μm, top 12%), a V1 neuron that receives retinal afferents from these local ON and OFF RGCs has a receptive field of weakly overlapping ON and OFF sub-regions. This results in a high simpleness index (SI, 0.42, see Methods), representing simple cell-like segregation between ON/OFF subregions. In contrast, when the distance between ON and OFF RGCs is small (Fig. 2c, purple circle, *d*_*ON-OFF*_ = 23 μm, bottom 5%), the inputs to V1 generate the receptive field of highly overlapping ON and OFF sub-regions with low SI (0.15), like a complex cell. In this scenario, the simple/complex tuning in V1 is simply destined from variation of the local arrangement of ON and OFF RGC mosaics.

Our model showed that variation of the response modulation ratio can be determined by the distance between ON and OFF subregions of RF. This involves *F*_*1*_*/F*_*0*_, the ratio of the first harmonic component to the mean elevation of the neuronal response to a drifting sinusoidal grating stimulus, another indicator of simple (*F*_*1*_*/F*_*0*_ > 1) or complex (*F*_*1*_*/F*_*0*_ < 1) cells. Based on the spike rectification model^20,25^, a nonlinear sigmoidal transfer function between the ON-OFF distance of RF and the *F*_*1*_*/F*_*0*_ of neural response was obtained analytically (Fig. 2d, see Supplementary Information and Supplementary Fig. 1 for details). From this result, we confirmed that the unimodal distribution of the ON-OFF distance shown in Fig. 2b can generate bimodal segregation of *F*_*1*_*/F*_*0*_ observed in the data (Fig. 2e).

### Periodic spatial organization of simple/complex cells

One important prediction arises from the result above: the spatial organization of simple and complex cells across a cortical layer would reflect the spatial layout of the *d*_*ON-OFF*_ in the RGC mosaic, organization into topographical clusters. As shown in Fig. 2f, the spatial distribution of *d*_*ON-OFF*_ is clustered across the RGC mosaics (Fig. 2f, top), generating local regions of large or small *d*_*ON-OFF*_ values. According to our model, it is predicted that simple and complex cells in V1 must appear as an organization of clusters across the cortical surface (Fig. 2f, bottom).

To test this idea, spatial organization of simple and complex cells in V1 was examined using published receptive field data^32^ obtained by multielectrode recording in cats (Fig. 3a). From the observed ON and OFF receptive fields, the simple/complex tuning index (SI) and the distance between ON/OFF center of mass (*d*_*ON-OFF*_) of each recording site were measured (Fig. 3b). The recording data contained both simple- and complex-like receptive fields, which showed segregated (left) or overlapped (right) ON and OFF sub-regions respectively. As reported^32^, the distribution of orientation preference varied periodically (Fig. 3c, top). Interestingly, both the spatial variation of *d*_*ON-OFF*_ and SI in V1 appeared periodically clustered along the cortical penetrations. The distribution of *d*_*ON-OFF*_ (Fig. 3c, 3^rd^ row) was well-fitted to a sinusoidal function of ∼1.1 mm period. Comparable to this, the distribution of SI (Fig. 3c, bottom) was also fitted to a sinusoidal function of nearly identical spatial period (∼1.0 mm) and phase (phase difference ∼17°). We found that the value of the observed SI and *d*_*ON-OFF*_ was tightly correlated as predicted by the model (n = 52 data points from 2 penetrations, Pearson correlation coefficient, *r* = 0.66, *p* = 1.2×10^−7^).

**Figure 3.**
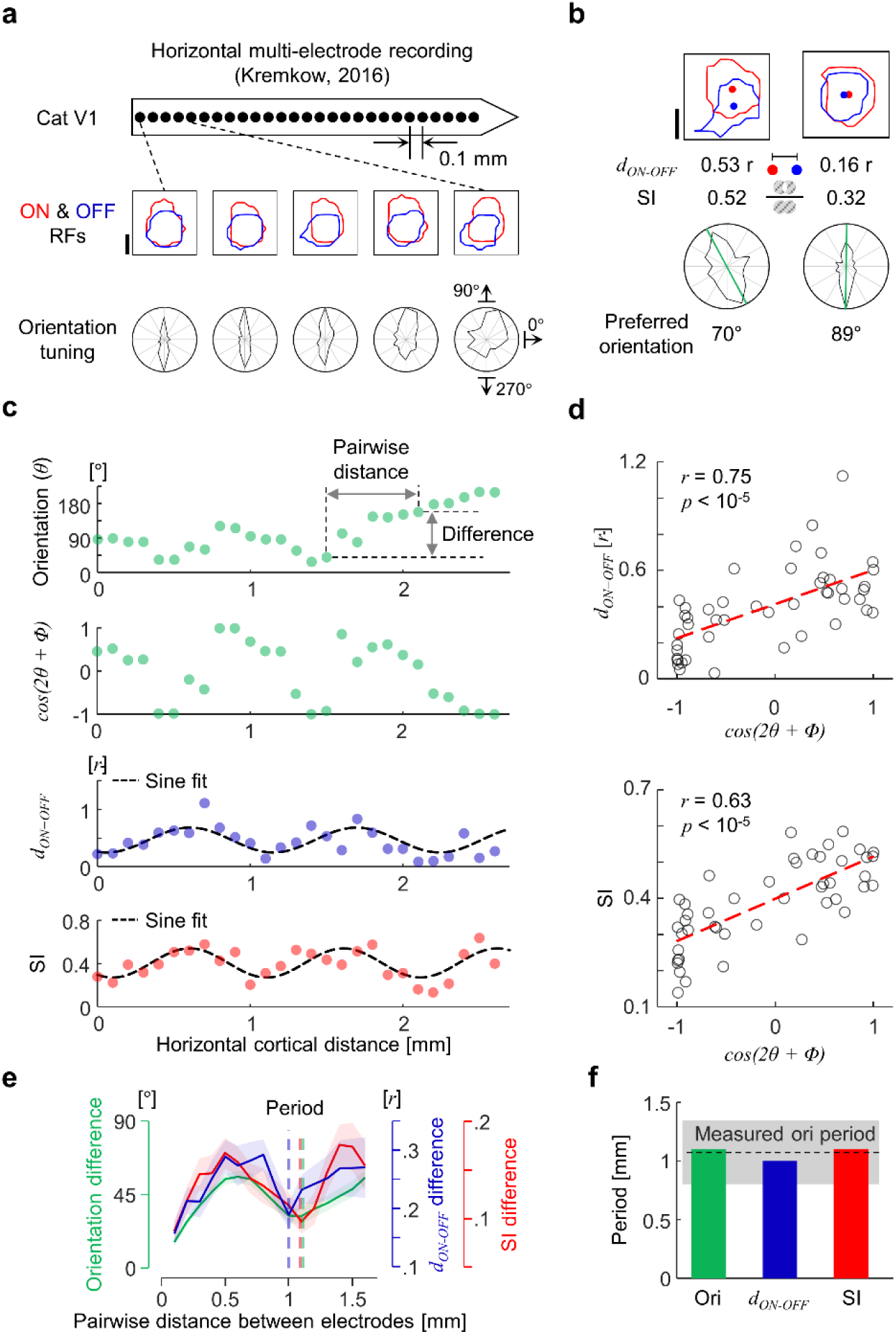
Periodic spatial organization of simple/complex cells and the common period with orientation preference in cat V1. (a) Illustration of multielectrode recording from Kremkow, 2016^32^. The contour of ON and OFF receptive fields (measured with light and dark stimuli, respectively) is defined as a level of z-score = 1.5 for each ON/OFF receptive field. Scale bar: average radius (*r*) of the ON/OFF receptive field for each penetration. Polar plots represent a normalized response to drifting bars. (b) Example calculation of *d*_*ON-OFF*_ (distance between center of mass of ON/OFF sub-regions), SI, and preferred orientation. (c) Periodic spatial clustering of orientation preference (green, and its phase-adjusted cosine values, Supplementary Figure 3), *d*_*ON-OFF*_ (blue), and SI (red). The Black dashed lines are sine fits for *d*_*ON-OFF*_ and SI. The *d*_*ON-OFF*_ fit: 0.47 + 0.22 × sin(*2πx/λ* - 1.92), *λ* = 1.1 mm (*R*^*2*^ = 0.41, *p* = 3×10^−4^). The SI fit: 0.4 *+* 0.13 × sin(*2πx/λ* - 2.19), *λ* = 1.0 mm (*R*^*2*^ = 0.52, *p* = 2×10^−5^). Phase difference between two fits: 17°. (d) Correlation between cosine of orientation (cos(*2θ+Φ*)) and *d*_*ON-OFF*_ (top), and SI (bottom) (2 penetrations, N = 52 sites). (e) Average pairwise difference as a function of pairwise distance (averaged over 2 penetrations, averaged pairwise sample > 20 for each pairwise distance). Dashed vertical lines represent the period of each curve (∼1.1 mm). (f) Comparison of the observed period of SI, *d*_*ON-OFF*_, and orientation with reported values of the orientation map periods^38^.

More interestingly, the spatial organization of *d*_*ON-OFF*_ and SI were correlated with orientation preference, and had a common period identical to that of the orientation tuning. For direct comparison, orientation preference (*θ*) was transformed into cos(*2θ+Φ*) (Fig. 3c, 2^nd^ row), and was shifted to find the maximum correlation (Supplementary Fig. 3). As shown in Figure 3d, the values of *d*_*ON-OFF*_ (or SI) and the cosine of orientation preference were correlated across cortical surface (Fig. 3d, n = 46 data points from 2 penetrations). Furthermore, the remarkably similar clustering period among the three organizations was manifested in the average absolute pairwise difference for each measure (orientation, *d*_*ON-OFF*_, and SI) plotted as a function of cortical distance (Fig. 3e, averaged over the 2-penetration data sets, where each mean value includes more than 20 pairwise comparisons). The calculated mean period values (orientation 1.1 mm, *d*_*ON-OFF*_ 1.0 mm, SI 1.1 mm) were not only similar to each other, but also matched the previously reported values of period of orientation maps in cats^38^ (Fig. 3f).

### Periodic clustering of simple/complex neurons from RGC mosaics

The observed periodic organization of simple/complex neurons, and their consistent period with the orientation preference, suggest that a common organizing principle may exist for tiling of both simple and complex tuning of neurons and their orientation tuning. Previously, theoretical studies suggested that topographic organization of various neural tunings may arise commonly from the spatial organization of RGC mosaics^26,28,33–35^ and recent observations reported that cortical orientation preference can be predicted by the spatial arrangement of ON and OFF afferents^30,32^, providing evidence for retinal origin of the cortical tunings. In addition to these findings, here we show that the observed clustering of simple/complex tuning is predicted and explained by the retinal development model proposed by Paik and Ringach^26,27^. In this model, two noisy hexagonal lattices of ON and OFF RGC mosaics generate a periodic interference pattern of a local ON-OFF dipole-like arrangement, called a moiré interference pattern (Fig. 4a, top). In this interference pattern, the ON-OFF distance and ON-OFF dipole angle changes periodically across the mosaics, with their spatial period denoted as *λ*_*m*_. As suggested in previous model studies^34,35^, we assumed that the response of a local V1 neuron is constrained by the structure of ON/OFF afferents from the RGC mosaics (Fig. 4a, bottom). In this scenario, orientation tuning is determined by the alignment angle of the ON and OFF RGCs, and the SI of a V1 neuron is determined by the segregation between ON and OFF RGCs of corresponding afferents.

**Figure 4.**
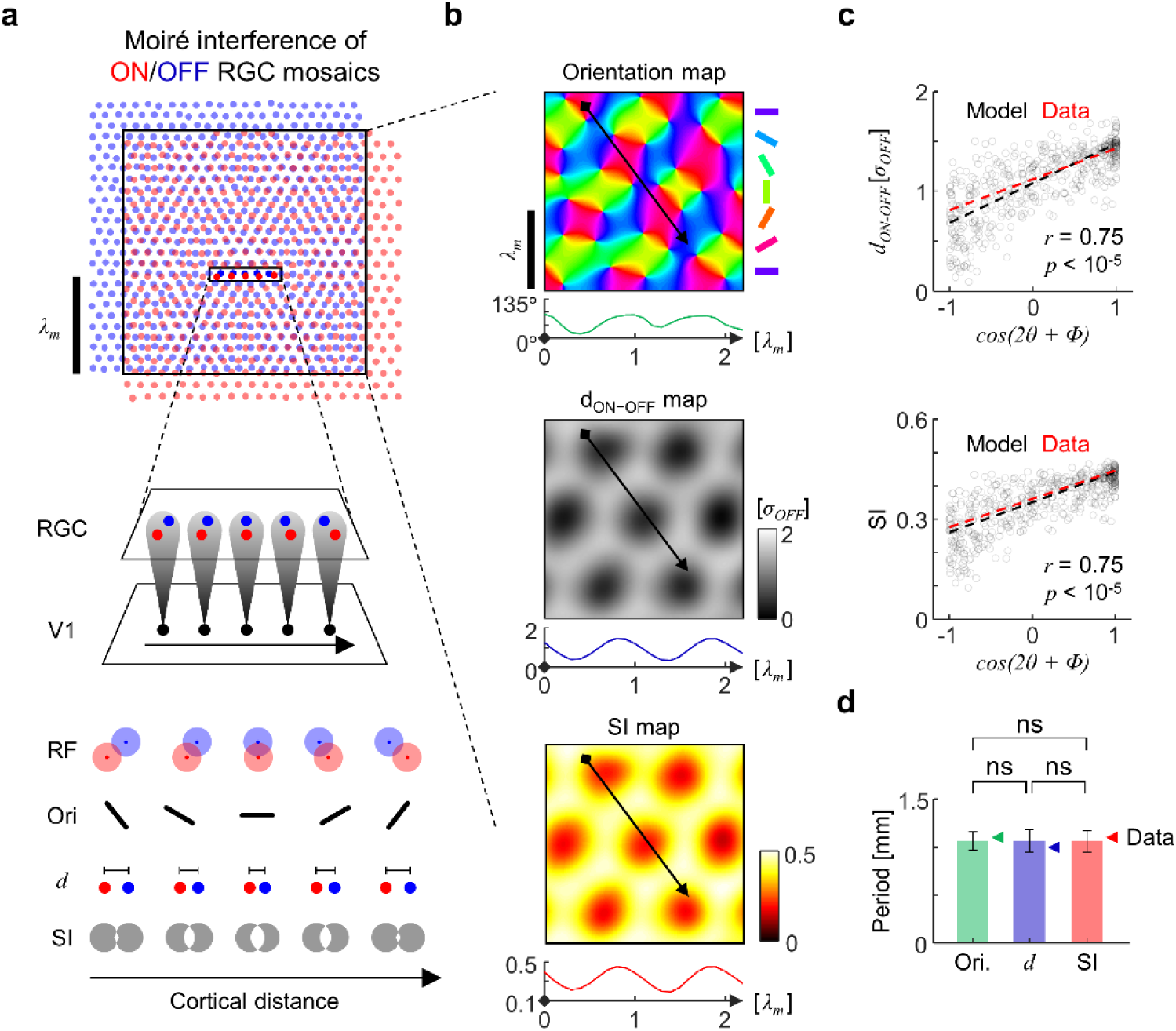
Moiré interference of retinal mosaics predicts the periodic spatial organization of SI, *d*_*ON-OFF*_, and orientation preference in V1. (a) ON (red) and OFF-center (blue) retinal ganglion cell receptive field mosaics are described as two noisy hexagonal lattices with different periodicity and the same angle. The resulting hexagonal moiré interference pattern has a characteristic period *λ*_*m*_, and can be described by local ON-OFF dipoles. Five example ON-OFF dipoles for ideal hexagonal moiré interference patterns occur in the region highlighted within a box (middle). Constructed receptive fields SI, *d*_*ON-OFF*_, and orientation preference of cortical neurons are represented (bottom) (b) Pseudo color representation of the synthetic map of preferred orientation, *d*_*ON-OFF*_, and SI (with example 1-D profiles along black arrow). *σ*_*OFF*_ represents the RF size of model OFF RGCs. (c) Correlation between cosine of orientation (*cos(2θ+Φ)*) and *d*_*ON-OFF*_ (top), and SI (bottom) obtained from diverse cortical penetrations on model maps (N = 100 penetrations). The data was rescaled to match the mean and standard deviation of the model for comparison. (f) Consistent periods obtained from three maps (one period value for each penetration for each map, N = 100 penetrations). ns: not significant (Wilcoxon rank-sum test, *p* = 0.25, 0.83, 0.13, for SI - orientation, SI - *d*_*ON-OFF*_, *d*_*ON-OFF*_ - orientation, respectively).

The model predicts that the preferred orientation, *d*_*ON-OFF*_, and SI, of V1 neurons are organized into a spatial cluster of the same period, *λ*_*m*_, and our model simulation results support this prediction (Fig. 4b, see methods for details). All three of the simulated maps showed clear periodic clustering of tuning across the cortical surface, matching the periodic organization of the RGC interference pattern. As in the data, cortical profiles of *d*_*ON-OFF*_ and SI in the model showed strong correlation with the cosine of the orientation preference (Fig. 4c). Furthermore, the period of each map, calculated from average pairwise difference as a function of pairwise distance, was identical to the predicted period *λ*_*m*_ of the retinal moiré interference (Supplementary Fig. 4). The distribution of the obtained period values of orientation, *d*_*ON-OFF*_, and SI from different locations of the model map were statistically indistinguishable from each other (Fig. 4d). These results imply that our retinal development model could explain the origin of correlated clustering of orientation preference and simple/complex tuning in V1.

### Prediction of local simple/complex tuning from information on local orientation tuning

Extension of the previous analysis of a correlated organization of simple/complex and orientation tuning motivated us to ask whether information on the local structure of an orientation map could predict the local simple/complex tuning of neurons in the corresponding local area. From the simulated maps of orientation and simple/complex tuning from a set of retinal mosaics, our model predicted that the locations of pinwheels are likely to be either maxima or minima of local simple/complex tuning, or SI (Fig. 5a). A previous study^27^ reported that two types of pinwheels of opposite polarity are generated from two distinct types of singularity of retinal interference patterns (see Fig. 1 in Paik and Ringach, 2012, for details). According to our model, these two singularities match the locations where ON-OFF RGC distance is either maximal or minimal, respectively. Thus, the model predicts spatial overlap of pinwheels of orientation maps and clusters of simple/complex cells: that is; the SI measured at each pinwheel location must be significantly higher or lower than that measured at other random locations.

**Figure 5.**
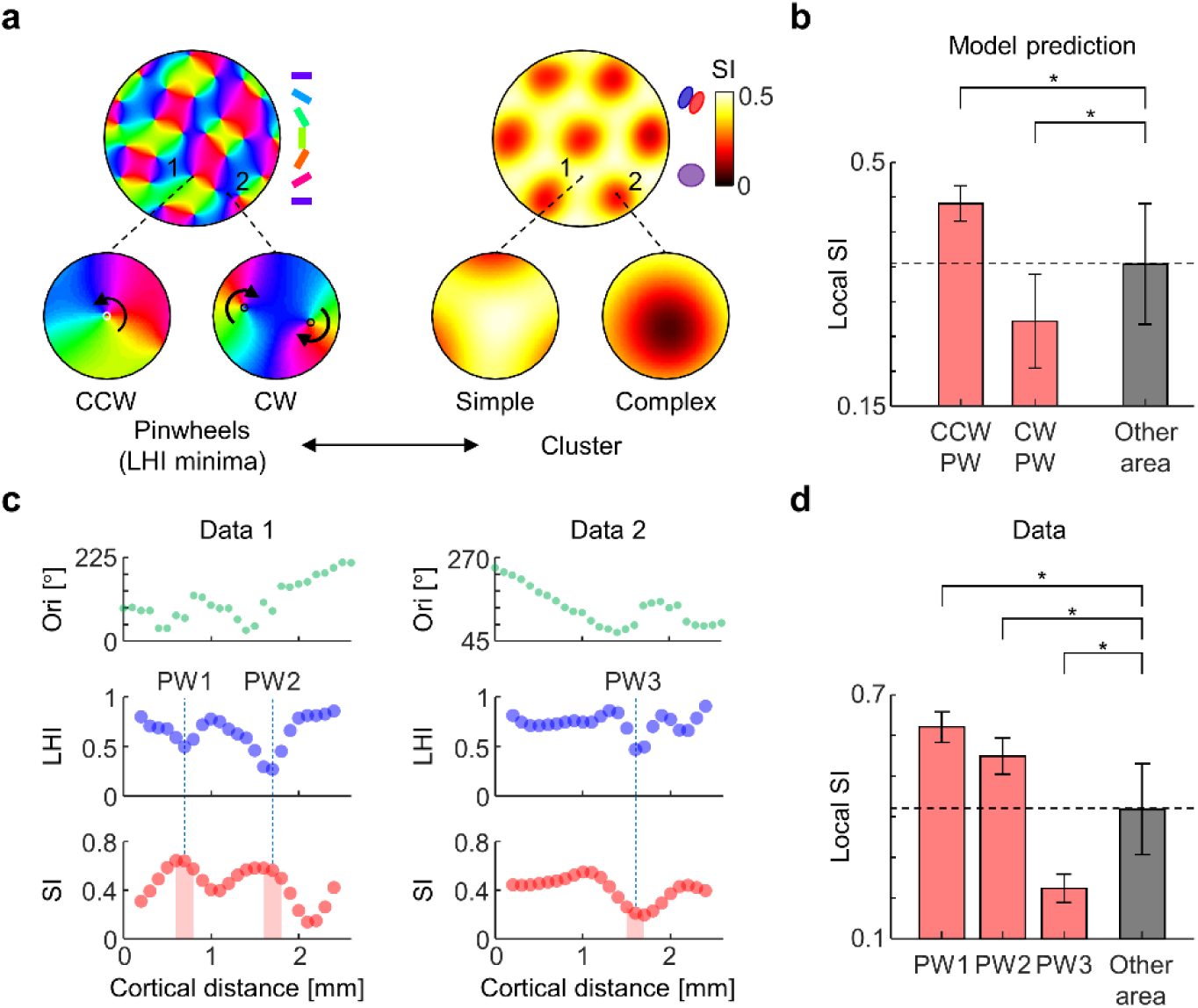
Inferring local simple/complex properties with local information of orientation map topography. (a) Comparison of topography of orientation map and SI map. (Left) Example pinwheels of the model orientation map with opposite winding polarity, counterclockwise (CCW, marked as 1) and clockwise (CW, marked as 2). (Right) Example SI of the model SI map at the corresponding locations (1 and 2). (b) Average SI over circular area within 1/8 map period from the center of simple and complex pinwheels (n = 179 simple pinwheels, n = 214 complex pinwheels) are significantly higher/lower than average SI over non-pinwheel areas (Wilcoxon rank-sum test, \**p* < 0.0001). (c) Orientation (green), LHI (blue) and SI (red, Gaussian-smoothed) in cat V1 recording data in Fig. 3 (2 penetrations). Three local minima of LHI (dashed black lines, putative pinwheels) and SI values near corresponding locations (red area) were identified. (d) Local SI near pinwheel locations in (c). Average SI were calculated for 100 µm range for each LHI minima. (Wilcoxon rank-sum test, \**p* = 0.0065, p = 0.027, p = 0.016 for PW1, PW2, and PW3, respectively).

To quantify this model prediction, we measured pinwheel locations on the simulated orientation maps, using the local homogeneity index (LHI, see Methods) minimized near pinwheels^39,40^. Then, SI values were measured near the center of each type of pinwheel (circular area within 1/8 map period). Those averaged SI values at each type of pinwheel were significantly higher or lower than the average SI at other random locations (Fig. 5b, \**p* < 0.0001, Wilcoxon rank-sum test). Thus, the model predicts “simple pinwheels” and “complex pinwheels” where SI values are local maxima or minima, respectively.

As the model predicted, we found evidence in the data that SI values are either maxima or minima at pinwheel locations. Similar to the model analysis above, 1-D LHI profiles were obtained from the orientation preference of recording data (Fig. 5c, from 2 penetrations). Although identification of the pinwheel polarity was not possible in this dataset due to the dimensions of the recordings, we found that the data contained three cortical locations where LHI values were locally minimized, implying that the recording passed near pinwheels. We identified these three locations of minimal LHI as tentative pinwheel locations (black arrows, PW1, PW2, and PW3) and found that the detected local maxima or minima of SI (triangles) were located close (within 100 μm) to the three tentative pinwheel locations. To quantify further this correlated architecture, we calculated average SI values of neighboring electrodes (within 100 μm) at each LHI minimum. The local SI values near pinwheels (LHI minima) were significantly higher (PW1, PW2) or lower (PW3) than those at other recording sites (Fig. 5d, red bars, \**p* = 0.0065, 0.027, and 0.016 for PW1, PW2, and PW3, Wilcoxon rank-sum test). This result suggests that our model can exploit the information of orientation map topography to predict the local simple/complex tunings, further supporting the validity of the model.

### Discussion

Our findings suggest that simple and complex tuning in V1 can commonly originate from spatial arrangement of the local projection of retinal afferents. Analysis of multielectrode recording data from cats revealed that simple and complex cells are periodically clustered in V1. This is the first report that simple and complex tunings are topographically organized in V1. Moreover, the observed periodic organization has a period consistent with the orientation preference, implying the common origin of the simple/complex cells and the orientation tuning in V1. The Paik-Ringach model predicts and explains the essential features of the observed periodicity from the periodic projection of retinal afferents imprinted in retinal mosaics. We further demonstrated that local orientation map topography and the local simple/complex tuning properties are correlated in a manner consistent with the model prediction.

In addition to afferent circuits, intracortical circuits can also contribute to the modulation in the simple/complex tuning within V1 as suggested in previous studies^19,21,22^. Indeed, our results do not rule out the possible role of intracortical activities after the initial tuning is constrained by the afferent inputs. However, from the observation that several experimental actions, such as silencing intracortical activity, did not change the orientation selectivity of V1 neurons due to thalamic inputs^41–43^, and that the arrangement of thalamic inputs can predict diverse tuning properties^30,32^, it is reasonable to suggest that the effect of intracortical inputs would be less influential to the initial development of simple and complex tuning than the effect of afferent inputs. Moreover, our results regarding the periodic spatial organization of simple and complex cells were predicted by the retinal afferent model^26^, which strengthens the view that major tuning properties of V1 neurons are anchored by retinal afferents, and that intracortical circuits refine or diversify a degree of tuning that the afferent circuits cannot solely develop^43^. The correlated architecture of simple/complex properties and the orientation map suggests further systematic combination of simple/complex tuning and other functional tunings in the thalamo-recipient layer of V1. Our model simulation and data analysis revealed that variation of simple/complex tuning is systematically tiled in relation to the underlying geometry of the orientation tuning, implying that the various feature selectivities in V1 are efficiently combined via systematic rules between functional maps^39,44–47^.

One might argue that the ratio between simple and complex cells in layer 4 is not consistent but instead is fairly different across species. For example, layer 4 of cat V1 is more dominated by simple cells^9^, while more complex cells are observed than simple cells in monkey V1^16^. Because a simple/complex tuning index could be modulated by intracortical activity^22^ or cortical nonlinearity^25^, difference of such parameters across species could elicit shifting of the distribution. More importantly, however, the tendency that both simple and complex cells develop simultaneously in the earliest hierarchy of visual cortex is commonly observed, which is consistent with our model prediction of retinal origin of simple and complex cells in V1. The results of several studies suggest that simple cells are mostly observed in the earliest layer of V1^48,49^, and that the proportion of complex cells becomes greater as the layers get deeper^9^. These observations are not different from our model prediction, because projection from an input layer of V1, especially layer 4, will converge into a superficial layer, such as layer 2/3, to generate more complex receptive fields. Rather, our finding suggests that the architecture of V1 is not only hierarchical but also parallel, and this parallel architecture refines the classical notion of visual cortex. That is, the role of simple/complex cells in visual information processing is not restricted to distinguishing different stages of the cortical microcircuits, but can be regarded as an element of functional diversity in the same cortical layer.

To sum up, the observed periodic spatial organization of simple and complex cells provides a population-level clue regarding how simple and complex receptive fields are generated and leads to the view that the distance between ON and OFF retinal afferents provides the source of the simple/complex spectrum. Complementary to the classical notion that simple and complex cells are hierarchically distinct, the observed periodic spatial organization of simple/complex cells shows systematic variation in the earliest layer in V1 that receives thalamic inputs. That the period is consistent with that of the orientation preference encourages the view that structured retinal afferents designed by interference between ON and OFF RGC mosaics provide the common source of both orientation preference and the simple/complex-property of the connected V1 neurons. These results support the theory that the diverse functional tunings of V1 are determined by the arrangement of ON/OFF afferent inputs of retinal origin.

## Methods

### Simpleness index (SI)

To quantify the simple/complex tuning of the receptive field, we calculated the simpleness index (SI), which represents the degree of segregation between ON/OFF subregions^5,17,30,50^. The SI is defined as follows:

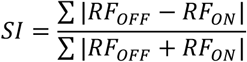

Where *RF*_*OFF*_ and *RF*_*ON*_ represent 2-d matrices of ON and OFF receptive field subregions, respectively, and the summation is over all matrix elements.

### Analysis of RGC mosaics

The ON-OFF dipole was defined as a line connecting the nearest ON cell from each OFF cell in the mosaic. The map of *d*_*ON-OFF*_ in Fig. 2f was obtained as,

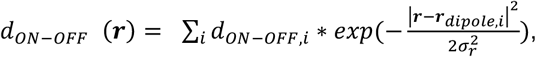

where ***r***_*dipole,i*_ and *d*_*ON-OFF,i*_ are the center and size of the *i*^*th*^ dipole, respectively (*σ*_*r*_ = 116 μm, average nearest distance between OFF cells). After the above calculation, the map was linearly rescaled to match the minimum and maximum value of the original *d*_*ON-OFF*_ values. The SI values in Figure 2 were calculated by assuming that a model V1 neuron receives inputs from one ON-center RGC and one OFF-center RGC of equal strength. The receptive field of ON- and OFF-center RGCs were modeled as the difference of a Gaussian (*σ*_*surround*_ = *3σ*_*center*_, *σ*_*center, ON/OFF*_ = half of the average ON-ON/OFF-OFF distance, respectively). The map of SI in Fig. 2f was obtained as the same formula for *d*_*ON-OFF*_.

### Receptive field data

Spatial organization of simple/complex cells in V1 was analyzed using published receptive field data obtained by multielectrode recording in Layer 4 of cat V1. This was provided by Jose-Manuel Alonso via data presented in Figure 2 of a previous study^32^. The detailed experimental procedures for mapping receptive fields are described in the reference.

We defined the size of the receptive field (*r*) for each recording as the average radius of receptive fields within each penetration (assuming circular receptive fields). The distance between the center of mass of ON and OFF subfields were normalized by dividing that distance by *r*. The period of each distribution (SI, *d*_*ON-OFF*_, and orientation preference) was calculated as the distance at which the pairwise difference value (Fig. 3f) reaches its minimum among local minima of the curves, following the process to calculate the period of orientation preference in the reference^32^.

### Analysis of homogeneity of the organization of orientation preference

To quantify the degree of homogeneity of the organization of orientation preference at a specific location *x*_*i*_, we calculated the local homogeneity index (LHI) with window size *σ* (170 μm) as follows^40^:

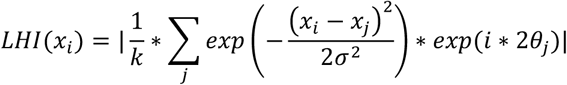

where *j* represents each site in the penetration, *k* is a normalization constant that makes the theoretical maximum value of LHI = 1, and *θ*_*j*_ is the preferred orientation of the *j*^*th*^ site. To avoid the edge effect in calculating the LHI, two sites at either end of the recordings were not represented. To obtain the LHI of the model orientation map, the same formula with the same window size (the sizes of model and data were normalized to match the period) was applied to two-dimensional space^39^. The location of pinwheels was identified as local minima of LHI that were smaller than the quartile.

### Map simulation

The simulations were conducted based on the statistical wiring model published earlier^26,34,35^. Here, we summarize the algorithm and parameters that were used to produce the results.

### Generation of retinal ganglion cell mosaics

The ON and OFF RGC mosaics used in the simulation were generated by adding random spatial noise to each node of the hexagonal lattices that represent the position of the center of ON-center and OFF-center receptive fields, respectively. The position vectors of the centers of the receptive fields were defined as

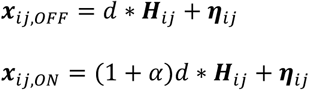

where *d* represents the lattice constant of the OFF mosaic, *(1+α)d* represents the lattice constant of the ON mosaic (*α* = 1/7), ***η***_*ij*_ represents the 2-D additive Gaussian noise with a standard deviation *σ* (= 0.05d), and ***H***_*ij*_ represents the vectors of the nodes of a unit hexagonal lattice spanned by two basis vectors.

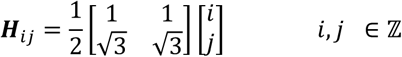

The characteristic period of the hexagonal moiré interference pattern, *λ*_*m*_, is given by^51^

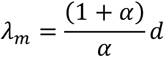

when the directions of the two principle axes of the lattices match.

The main results of the model simulation and comparison with the data is nearly identical to the various choices of the parameter of the moiré interference.

### Statistical connectivity and receptive field computation

The mean receptive field at each cortical site can be computed by the weighted sum of the afferent LGN input (it relays the afferent RGC input).

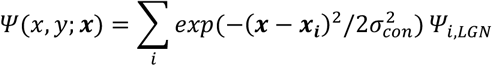

where ***x*** is the cortical site at which we calculate the mean receptive field, ***x***_*i*_ is the location of the *i*^*th*^ LGN afferent, *Ψ*_*i,LGN*_ is the receptive field of the LGN afferent, and *σ*_*con*_ (= 0.28d) is the parameter that determines the spatial extent of the synaptic weight distribution, which is assumed to be a form of Gaussian^35^.

### Measurements of cortical maps

Simulated cortical maps were obtained from the computed receptive fields at each cortical position. The SI of the V1 neurons was calculated in the same way as the SI of the data. The preferred orientation of each receptive field was calculated as the angle orthogonal to the line connecting the center of ON and OFF subregions. If either an ON or OFF subregion dominated (so called “monocontrast” cells, which respond to only one particular sign of contrast), so that the sum of all the weights of ON afferents were larger than two times the sum of all the weights of OFF afferents and vice versa, the neurons were excluded from the map measurement. After obtaining the SI and orientation preference of each cortical site, we smoothened the map with a 2-D Gaussian kernel with standard deviation 0.16 *λ*_*m*_. The filtered map of SI and *d*_*ON-OFF*_ were linearly rescaled to recover the minimum and maximum values of the raw cortical maps.

To compare the data and model, we obtained pixel-values from cross-sections of each simulated map along line segments that had the same length as the data segments, but with random penetration direction. The length of the data and model was normalized to match the period of orientation preference of the data (1.1 mm) and that of model (*λ*_*m*_). As in the data, 27 sites with equal spacing were sampled for each cross-section (10,000 cross-sections) and pairwise difference curves were calculated. Two pairwise difference curves were randomly sampled and the mean and standard deviation of the mean of the two curves were calculated for 100,000 iterations. For pairwise differences of SI and *d*_*ON-OFF*_, the scale of variation was normalized by dividing the maximum value of the mean model curve and multiplying by the maximum value of the data curve.

## Acknowledgements

We are grateful to Jose-Manuel Alonso (State University of New York) for sharing receptive field data on the cat primary visual cortex, via data presented in Figure 2 of his reference^32^. This work was supported by the National Research Foundation of Korea (NRF) grant funded by the Korea government (MSIT) (No. NRF-2019R1A2C4069863, NRF-2019M3E5D2A01058328) (to S.P.).

## Supplementary Information

### Model of the relationship between the membrane voltage and *F*_*1*_*/F*_*0*_ ratio

Following previous modeling studies^20,25^, the membrane voltage response of a V1 neuron to the drifting grating stimulus can be expressed as a sinusoidal function:

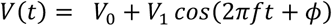

where *V*_*0*_ represents the mean elevation of the membrane voltage and *V*_*1*_ represents the amplitude of the modulation, *f* is the temporal frequency of the drifting grating, and *ϕ* si a constant phase term.

The relationship between the membrane voltage fluctuation and *F*_*1*_*/F*_*0*_ was formulated using the following 3-parameter model:

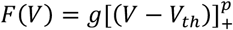

where *F* is spike rate, *p* is an exponent and *g* is a gain factor. For the simplest case, *p* = 1, the analytic expression of *F*_*1*_*/F*_*0*_ as a function of a variable *χ* = (*V*_*th*_ − *V*_0_)/*VV*_1_ can be obtained as follows^20^.

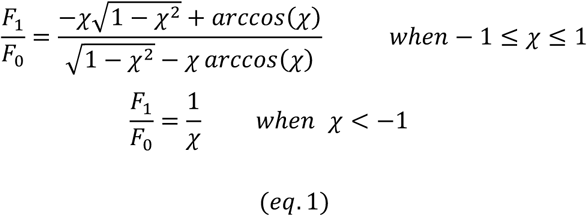

For a fixed value of *V*_*th*_, *F*_*1*_*/F*_*0*_ can be described as a function of variable *V*_*1*_*/V*_*0*_.

The work of Mechler and Ringach^20^ suggests that such a nonlinear relationship can induce the bimodal distribution of *F*_*1*_*/F*_*0*_ even if the underlying distribution of *χ* (or *V*_*1*_*/V*_*0*_) is u nimodal. T hus, s imple (*F*_*1*_*/F*_*0*_ > 1) and complex (*F*_*1*_*/F*_*0*_ < 0) cells can be considered as a common type of cells. However, what factor in visual circuit can make the spectrum of such variables was not fully understood.

### Relationship between ON-OFF RGC afferent distance and *F*_*1*_*/F*_*0*_ of V1 neuron

Here, we advance the notion of the previous studies by suggesting that the distance between ON and OFF RGC afferents can give the source of the spectrum of *F*_*1*_*/F*_*0*_. An analytically tractable model that expresses *F*_*1*_*/F*_*0*_ as a function of the distance between ON and OFF RGC afferents are demonstrated in this section.

A previous electrophysiological study showed that the response measured by firing of retinal ganglion cells varies sinusoidally with the matched temporal frequency of the drifting grating stimulus with optimal spatial frequency^52^. Thus, we start by writing a firing rate of an RGC to the drifting grating stimulus as a sinusoidal function, which is denoted as r(t).

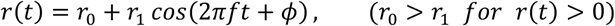

where *f* is the temporal frequency of the drifting grating stimulus and *ϕ* is the phase determined by the location of the RGC receptive field (*x*).

Note that for a suitable choice of reference, one can write the phase as a variable of spatial position divided by the spatial frequency (*λ*) of the drifting grating.

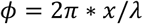

Here, the value of *λ* was determined to produce the maximum response for each RGC, where the RGC receptive field was modeled as in the main text and the response was calculated with a conventional linear nonlinear model^53^.

Summation of the response of ON and OFF RGCs at different positions (and thus different phases) yields the equation for the summed response, *r*_*sum*_(*t*) (Supplementary Fig. 1a, 1^st^ column).

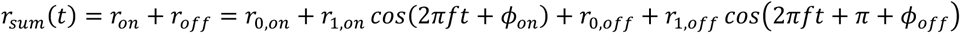

The addition of *π* phase arises from the opposite polarity of ON/OFF response. By letting *r*_0,*on*_ = *r*_0, *off*_ = *r*_0_, *r*_1, *on*_ = *r*_1, off_ = *r*_1_, and applying the trigonometric identity yields the simplified expression of the summed response.

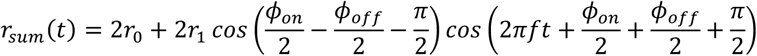

The interpretation of the above equation is as follows. The first term, 2*r*_0_, is independent of the phase difference between ON and OFF RGCs. The amplitude of the second term, however, is dependent on the phase difference between ON and OFF RGCs (*ϕ* _*on*_ − *ϕ*_*off*_). When *ϕ* _*on*_ = *ϕ* _*off*_, namely when ON and OFF receptive fields are completely overlapped, the amplitude of sinusoidal modulation becomes zero due to the 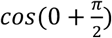 term. As *ϕ*_*on*_– *ϕ*_*off*_ increases, the amplitude of sinusoidal modulation increases and becomes maximum when *ϕ*_*on*_– *ϕ*_*off*_ = *π*.

Next, the expression of the membrane voltage fluctuation suggested by Mechler and Ringach was linked with the above expression of *r*_*sum*_(*t)* as follows (Supplementary Fig. 1a, second column).

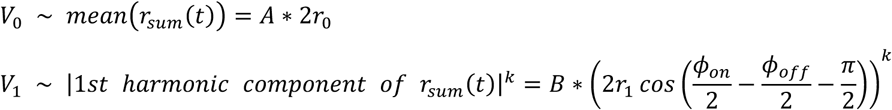

where the exponent *k* was applied for the expansive relationship between input and membrane voltage modulation that arises from nonlinear integration in dendrites^54^. This nonlinear relationship can generate a skewed distribution of *V*_*1*_*/V*_*0*_ as observed in cat^25^ from a Gaussian-like distribution of ON-OFF distance (Supplementary Figs. 1b, c).

This sinusoidal membrane voltage fluctuation is rectified to generate spike response modulation (Supplementary Fig. 1a, third column). The expression that links the distance between ON/OFF afferent and *χ*, which determines the modulation ratio *F*_*1*_*/F*_*0*_ becomes,

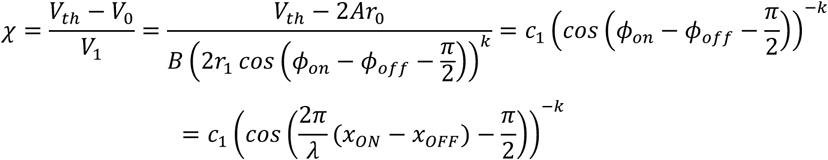

Where 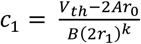 is a constant. Substituting the *χ* into eq. 1 and expressing the ON-OFF distance as *x*_*ON*_ – *x* _*OFF*_ = *d*, yields the analytic expression of *F*_*1*_*/F*_*0*_ in terms of *d*.

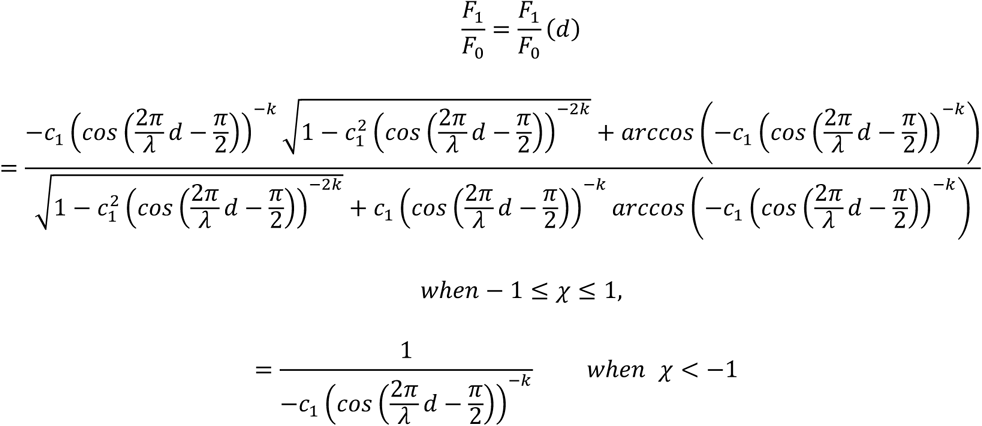

From this expression, the distribution of *F*_*1*_*/F*_*0*_ can be calculated as in the Fig. 2d of the main text. In our demonstration, *k* = 3, *c*_*1*_ *∼ N (μ, σ*^*2*^*) = N* (−0.04, 0.08^2^) were used (*N* represents the normal distribution).

---

**Supplementary Figure 1.**
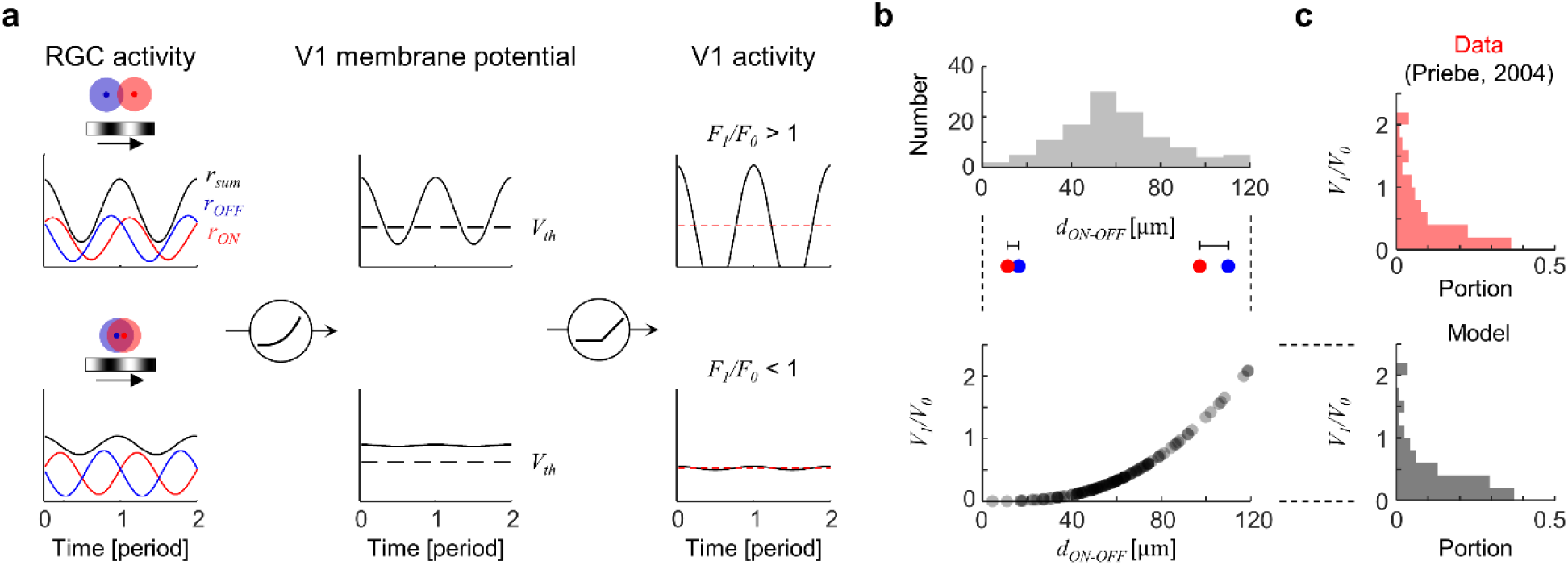
Relationship between *d*_*ON-OFF*_ and response modulation of V1 neuron to drifting grating. (a) Illustration of responses of RGCs and V1 neurons to the drifting grating stimulus following the model described in Supplementary Information. Dashed black line (*V*_*th*_) represents spike threshold. Dashed red line represents mean response. (b) Distribution of *d*_*ON-OFF*_ shown in main text (top) and model relationship between *V*_*1*_*/V*_*0*_ and *d*_*ON-OFF*_ (bottom). (c) Skewed distribution of *V*_*1*_*/V*_*0*_ in both top (adapted from Priebe, 2004^25^) and model (bottom, *B (2r*_*1*_)^*k*^ */ 2 Ar*_*0*_ = *7*: constant that linearly controls the scale of *V*_*1*_*/V*_*0*_).

**Supplementary Figure 2.**
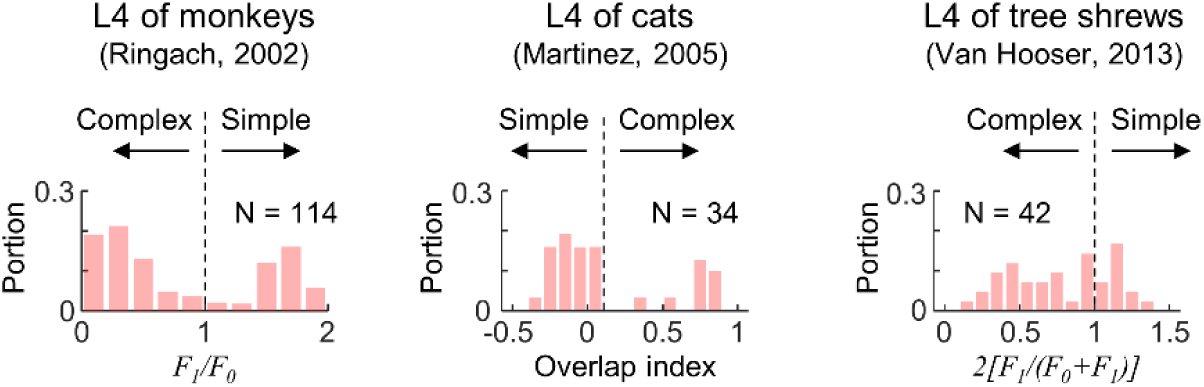
Distribution of simple and complex cells in monkeys^16^, cats^9^, and tree shrews^17^. Simple and complex cells coexist in the earliest stage of visual cortex, layer 4, in monkeys, cats, and tree shrews. *F*_*1*_*/F*_*0*_ and *2[F*_*1*_*/(F*_*0*_*+F*_*1*_)] represent the degree of response modulation to sinusoidal drifting gratings, and the overlap index measures the degree of overlap between ON and OFF subregions of the receptive fields.

**Supplementary Figure 3.**
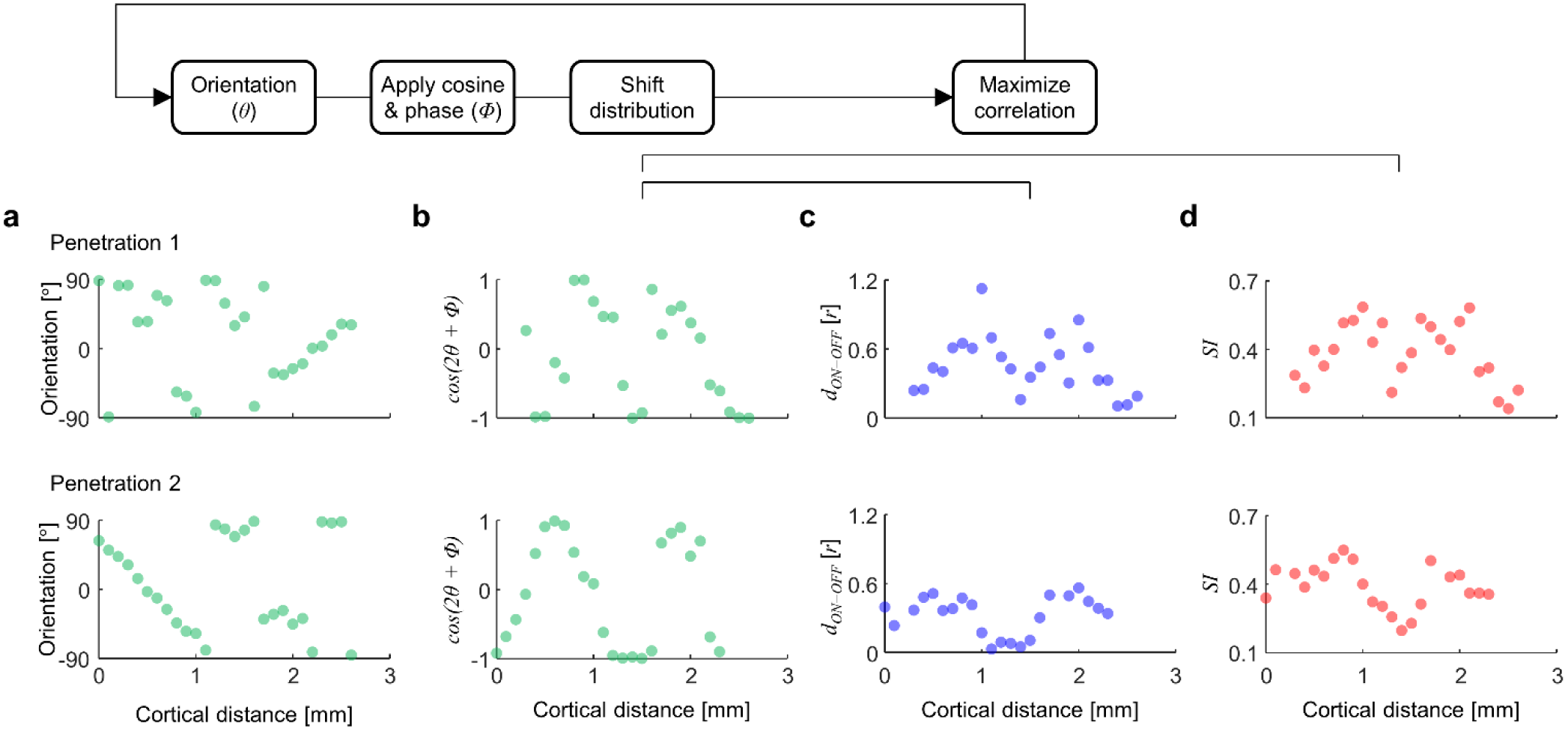
Quantifying correlation between orientation and SI (or *d*_*ON-OFF*_). (a) Spatial profiles of orientation preference (*θ*). (b) Transformed and distance-shifted profiles of *cos(2θ+Φ)*. (c) Shifted profiles of *d*_*ON-OFF*_ and SI. The values of *Φ* (0–360&) and shifting lag (−0.5–0.5 mm) were determined to maximize the correlation between (b) and (d). In position-shuffled control (shuffle position information of SI), the probability of obtaining as high a correlation value as in the data was significantly low (*p* = 0.005).

**Supplementary Figure 4.**
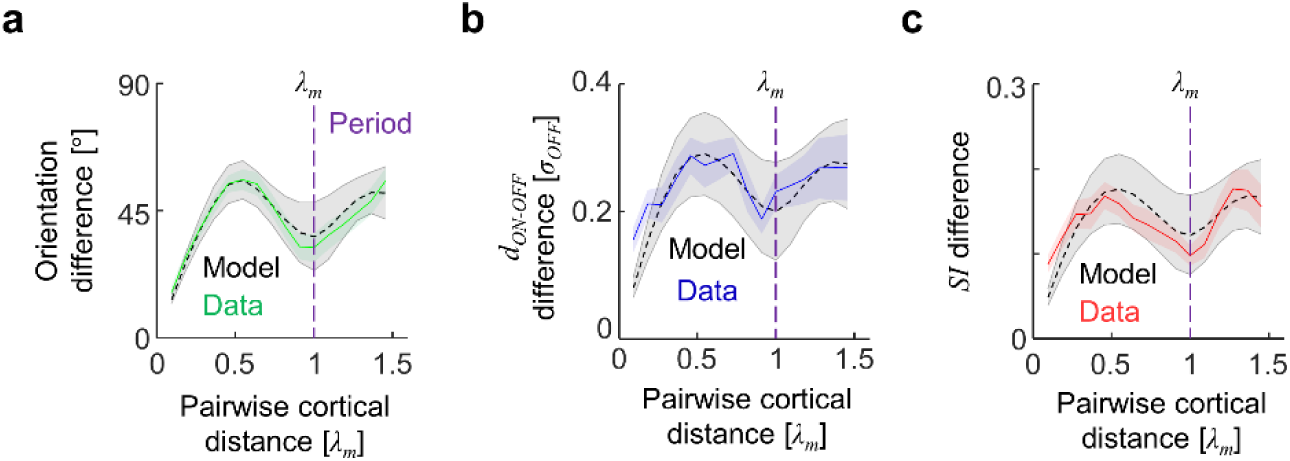
Common periods for orientation, *d*_*ON-OFF*_, and SI in model and data. Pairwise differences of (a) Orientation (b) *d*_*ON-OFF*_, and (c) SI for the simulated model maps. The common period *λ*_*m*_ is denoted as a purple dashed line. Shaded gray areas represent the standard deviation obtained from different cortical penetrations. For (b) and (c), the curves for data and model were normalized to match the maximum value.

